# Combining Ensemble Learning Techniques and G-Computation to Investigate Chemical Mixtures in Environmental Epidemiology Studies

**DOI:** 10.1101/147413

**Authors:** Youssef Oulhote, Marie-Abele Bind, Brent Coull, Chirag J Patel, Philippe Grandjean

## Abstract

**Background:** Although biomonitoring studies demonstrate that the general population experiences exposure to multiple chemicals, most environmental epidemiology studies consider each chemical separately when assessing adverse effects of environmental exposures. Hence, the critical need for novel approaches to handle multiple correlated exposures.

**Methods:** We propose a novel approach using the G-formula, a maximum likelihood-based substitution estimator, combined with an ensemble learning technique (i.e. SuperLearner) to infer causal effect estimates for a multi-pollutant mixture. We simulated four continuous outcomes from real data on 5 correlated exposures under four exposure-response relationships with increasing complexity and 500 replications. The first simulated exposure-response was generated as a linear function depending on two exposures; the second was based on a univariate nonlinear exposure-response relationship; the third was generated as a linear exposure-response relationship depending on two exposures and their interaction; the fourth simulation was based on a non-linear exposure-response relationship with an effect modification by sex and a linear relationship with a second exposure. We assessed the method based on its predictive performance (Minimum Square error [MSE]), its ability to detect the true predictors and interactions (i.e. false discovery proportion, sensitivity), and its bias. We compared the method with generalized linear and additive models, elastic net, random forests, and Extreme gradient boosting. Finally, we reconstructed the exposure-response relationships and developed a toolbox for interactions visualization using individual conditional expectations.

**Results:** The proposed method yielded the best average MSE across all the scenarios, and was therefore able to adapt to the true underlying structure of the data. The method succeeded to detect the true predictors and interactions, and was less biased in all the scenarios. Finally, we could correctly reconstruct the exposure-response relationships in all the simulations.

**Conclusions:** This is the first approach combining ensemble learning techniques and causal inference to unravel the effects of chemical mixtures and their interactions in epidemiological studies. Additional developments including high dimensional exposure data, and testing for detection of low to moderate associations will be carried out in future developments.

## Background

Biomonitoring studies of environmental chemicals demonstrate that the general population experiences exposure to multiple chemicals from many different sources and at varying levels. In addition to chemicals, non-chemical exposures such as drugs, infectious agents, macro- and micronutrients, and psychosocial stressors exhibit a significant role in the etiology of disease either independently or in combination with chemical stressors (1). The inception of the exposome concept (i.e. potentially including all exposures of potential health significance) (2), coupled with high throughput omics technologies, brings the field of environmental epidemiology to a new era, and large observational datasets will be increasingly used to address health effects of chemical exposures. However, implementation of the exposome concept is challenged by the ability to accurately assess the effects of multiple exposures (3).

The large majority of observational environmental epidemiology studies consider each chemical separately when assessing the potential adverse health effects of environmental exposures. Investigating single environmental exposures in isolation does not reflect the actual human exposure circumstances nor does it capture the multifactorial etiology of health and disease (4). Ignoring the potential confounding from correlated environmental exposures can lead to invalid conclusions, even if we consider that associational analyses may be valid for causal inference assuming that exposure data are accurate. These limitations have been recognized as critical by the National Institute of Environmental Health Sciences (NIEHS), as demonstrated by the inclusion of the topic of combined exposures in the NIEHS 2012-2017 strategic plan and the organization of two recent NIEHS workshops on chemical mixtures.

Several statistical methods have been proposed to estimate health effects of environmental mixtures, often with an emphasis on variable selection. These methods include Environmental-Wide Association Studies (EWAS) (5), penalized regression methods (e.g. least angle selection and shrinkage operator [LASSO]) (6), dimension reduction methods, and exposure-response surface methodology such as generalized additive models and kernel regression methods (7). In a recent study by Agier and colleagues used simulation to demonstrate feasibility of some established and emerging methods for handling chemical mixtures (8), including EWAS, Elastic net, sparse partial least squares, a “deletion/subtraction/addition” method (9), and Graphical Unit Evolutionary Stochastic Search (10). While promising, these methods underperformed when the goal was identification of individual exposures with an impact on the phenotype of interest. They tended to exhibit high false discovery proportions as the number of correlated exposures increased. Moreover, results from a recent NIEHS workshop showed that none of the tested approaches appeared to outperform the others (11).

Machine learning methods have a great potential for quantifying the role of chemical mixtures in regard to their effects on human health, in addition to their ability to detect complex interactions. Surprisingly, only a few studies have applied these methods in the context of environmental mixtures investigations (12, 13). Machine learning methods are mainly used for their excellent predictive ability. They typically consist of two steps, a first step where the algorithm “learns” the variables that are associated with the outcome, and a second step where the algorithm is tested in an independent dataset to estimate generalizability of the algorithm (14). Tree-based methods (15) and their variants, including random forests (16) and stochastic gradient boosting (17) are among the most popular machine learning method. These algorithms select combinations of variables, or exposures, which are predictive of the outcome and form a “rule”, or decision tree, based on these combinations. Rules or decision trees can thereby quantify the association between multiple exposure variables and a given outcome (14). Other newer machine learning algorithms include Bayesian Additive Regression Trees (BART (18)), Bootstrap aggregating (Bagging (19)), and Extreme Gradient Boosting (XGB). These algorithms can yield excellent predictive performance, and can provide Variable Importance Measures (VIMs) for the exposures. However, these measures are mainly based on predictive accuracy, and do not summarize the magnitude or direction of the association easily.

In this paper, we propose to use an ensemble machine learning technique called Super Learner (20) that offers greater flexibility in approximating the data generating mechanism, and combine it with G-computation (21, 22), a causal inference approach that can yield valid causal effect estimates. The proposed approach can mitigate the problems of multicollinearity and model misspecification, with nonparametric prediction algorithms fitting complex exposure-response curves. We extend the approach to reconstruct valid exposure-response relationships, and to infer estimates of potential two-way interactive effects.

## METHODS

### G-computation

G-computation is a maximum likelihood based substitution estimator of the G-formula (21). Application of this method allows using observational data to estimate parameters that would be obtained in a perfectly randomized controlled trial. Under the assumption that adjustment for observed confounders is sufficient to achieve independence between potential outcomes and the exposure levels, these estimates can be interpreted causally. Implementation of the G-computation estimator is equivalent to using the marginal distribution of the covariates as the standard in standardization, a familiar class of procedures in epidemiology (22). It is implemented by modeling the outcome as a function of the exposure and covariates. The fitted model is then used to predict the outcome under different exposure scenarios to be compared. The average causal effect (ACE) is therefore estimated by averaging the difference between the model predictions for all individuals across the desired exposure levels. Typically, g-computation relies on parametric models, however, in this novel approach; we use the SuperLearner ensemble learning technique to relax a priori assumptions about the underlying function since the nature of the exposure-outcome relationship is usually unknown.

### SuperLearner

SuperLearner is a data adaptive approach that has been proposed by van der Laan et al. (20, 23). It uses cross-validated risks to find an optimal combination of predictions from a list of algorithms supplied by the user that minimizes a given loss function (e.g. squared error). One of the most important properties of the SuperLearner is that it converges to the oracle estimator, i.e. it has been demonstrated that this convex combination performs asymptotically at least as well as the best choice among the library of candidate algorithms if the library does not contain a correctly specified parametric model, and it achieves the same rate of convergence as the correctly specified parametric model otherwise (24). The method therefore yields the closest approximation to the real data generating mechanism for a given set of candidate models. The weights applied to the convex combination are derived to minimize the prediction error.

### Simulation study

We used a matrix X of 5 exposure variables with a sample size n 300 from a Faroese birth cohort (25), where X = PCB, PFOS, *pp*'DDE, PFOA, HCB) represents concentrations of 5 weakly to highly correlated exposures (0 < *ρ* < 0.87), namely sum of polychlorinated biphenyls (PCB), Perfluorooctanesulfonic acid (PFOA), Dichlorodiphenyldichloroethylene (*pp’*DDE), Perfluorooctanoic acid (PFOA), and Hexachlorobenzene (HCB) (Figure 1). We also generated a binary variable for sex from a Bernoulli distribution, with a probability of being female of 50%. The 5 exposures were log2-transformed and standardized before simulating the exposure-response relationships to approximate a Gaussian distribution and to simplify interpretation.

**Figure 1:**
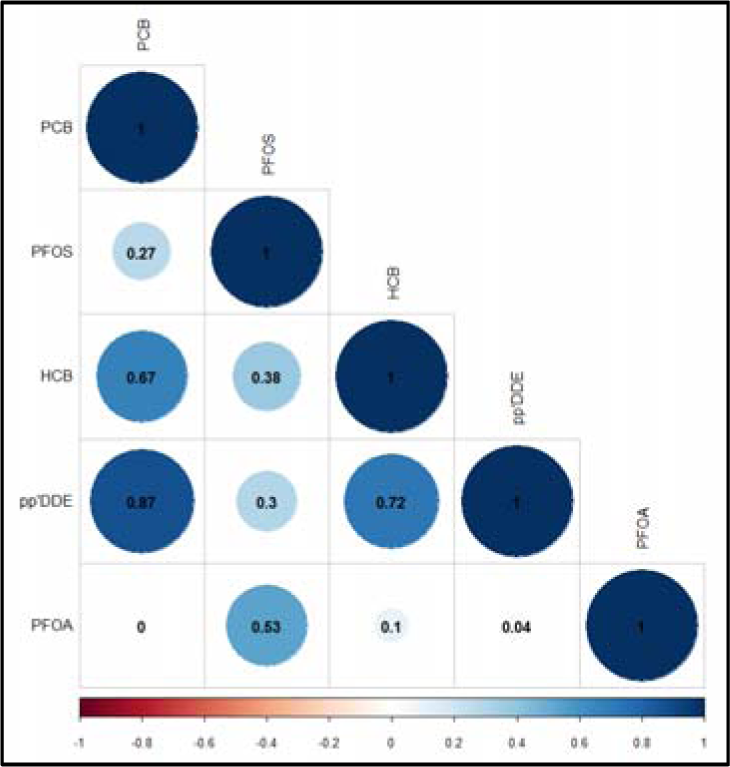
Correlation plot of the 5 environmental exposures

We generated four continuous outcomes with increasing complexity in regard to the exposure-response relationships. The properties of these simulated relationships and values of the *β* coefficients are described in table 1.

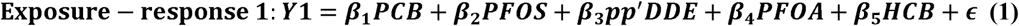

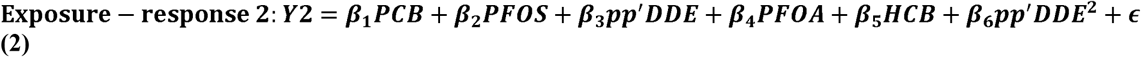

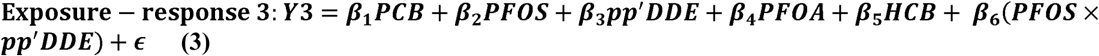

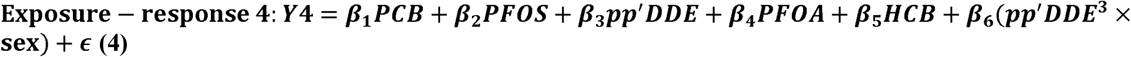

**Table 1:** Values of the coefficients for the 4 simulated exposure-response relationships.

| Coefficient | Variable | Simulation protocol ( values of $\beta$ coefficients) | | | |
| --- | --- | --- | --- | --- | --- |
|  |  | Y1 | Y2 | Y3 | Y4 |
| $\beta_1$ | PCB | -1 | 0 | 0 | 0 |
| $\beta_2$ | PFOS | 0 | 0 | -1 | -1 |
| $\beta_3$ | $pp'$DDE | -2 | -1 | -2 | -1 |
| $\beta_4$ | PFOA | 0 | 0 | 0 | 0 |
| $\beta_5$ | HCB | 0 | 0 | 0 | 0 |
| $\beta_6$ | $pp'$DDE<sup>2</sup> | - | -1 | - | - |
| $\beta_6$ | PFOS × $pp'$DDE | - | - | -3 | - |
| $\beta_6$ | $pp'$DDE<sup>3</sup> × sex | - | - | - | -2 |

### Analysis Method

For each data generating mechanism, we created 500 simulated datasets with similar exposure distributions to evaluate the predictive performance of the SuperLearner. To the data from each of the simulated exposure-response scenarios, we applied the SuperLearner algorithm. We included a convenient set of prediction algorithms in the library that can cover a large range of exposure-response relationships: the traditional generalized linear (GLM) and additive models (GAM), Elastic net regularization (26), multivariate adaptive polynomial spline regression (27), and Random Forests (16).

The primary measure of predictive performance was the Mean Squared Error (MSE), where lower MSE indicates a better estimate. For each algorithm included in the SuperLearner library, we present the 10-fold cross validated MSE and the *R^2^* for each of the scenarios with:

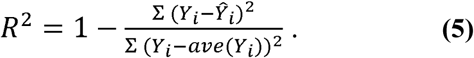

To compare the selected algorithms with the most commonly used method in the field, we also use the relative MSE (*rel*MSE) as a measure of relative performance to the widely employed linear regression as previously reported (28).

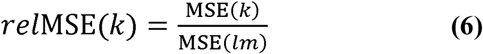

where k is the specific algorithm to assess (28). All these performance indices were averaged across the 500 datasets for each scenario.

#### Effect estimates using G-computation

For more complex ensemble machine learning techniques such as the SuperLearner, no simple parametric description that is comparable to the linear regression coefficients is available; we therefore use G-computation to infer valid effect estimates for specific exposures of interest. Let Y be the outcome and X = X_1_,…..,X_p_ a vector of predictors. Let’s also assume that we are interested in the effect estimates of a subset of environmental exposures, for each exposureX_j_, X_-j_ = (X_1_, X_2_ …..,X_j−1_, X_j+1_,……,X_p_) includes all the remaining covariates required for the identifiability of the effect estimate of X_j_. Each X_j_ can take values x ∈ τ, where τ is a continuous domain. The potential outcome under the exposure level X_j_ = x is denoted Y(x), whereas the potential outcome under the exposure level X_j_ = x + Δx is denoted Y(x + Δx). For example, in our simulation Δx = 1 corresponds to a one standard deviation (SD) increase in the log_2_-transformed values of exposure X_j_, and the parameter capturing the corresponding incremental change in the outcome Y can be expressed as follows:

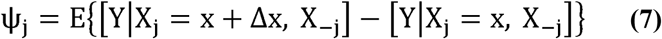

where the individual level effects are marginalized over the distribution of X_−j_. The estimator for the marginal effect of exposure X_j_ can be defined as follows:

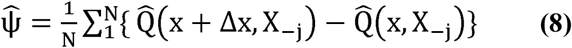

where 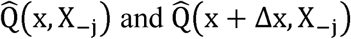 are the estimated potential outcomes for each individual under the exposures x and x + Δx, respectively, and N is the number of subjects in the sample. We derived these estimates non-parametrically using the SuperLearner by plugging in the exposure levels of interest for X_j_ and obtaining the predicted outcomes. In this simulation study, the average marginal effect for a one SD increase in a given log_2_-transformed exposure corresponds to the average difference between predictions from a dataset where the exposure of interest is replaced by 1(X_j_ = 1 and predictions from a dataset where the exposure is replaced by 0 (X_j_ = 0):

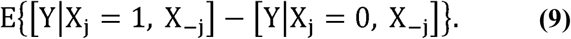

The results from the SuperLearner predictions were evaluated on the basis of False Discovery Proportion (FDP; the proportion of selected variables that were not truly related to the outcome), and sensitivity (the proportion of true predictors that were actually selected by the method). Additionally, we calculated the percent bias for each scenario.

#### Reconstructing the dose response relationship

In this proposed method, we use the SuperLearner-predicted outcomes at each specified level of the exposure of interest to construct a dose response relationship. In other words, we predict the outcome for all the unique values of X_j_ while keeping values of the variables in X_−j_ at their observed levels. Therefore, the average partial relationship between X_j_ and Y can be expressed as follows:

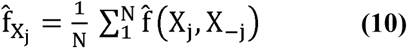

where N is the number of observations and 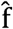 denotes predictions from the SuperLearner.

#### Detection of interactions

Suppose we are interested in detecting and estimating the effect attributable to the interaction between exposures X_1_ and X_2_ from a set of exposures of interest. After standardizing the exposures, the Interactive effect (IE) is expressed as:

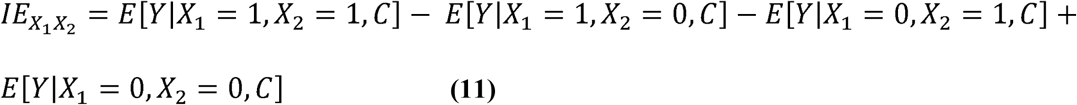

where C is a set of measured confounders sufficient to identify the effect estimates of *X*_1_ and *X*_2_. The formulations in this equation respectively correspond to the predicted outcome among those exposed to both *X*_1_ and *X*_2_, the predicted outcome for those exposed only to *X*_1_, the predicted outcome for those exposed only to *X*_2_, and the baseline outcome, i.e. predicted outcome for those exposed to neither *X*_1_ nor *X*_2_. Thus, if for example the result from the formula is significantly > 0, there is evidence that *X*_1_ and *X*_2_ work in a synergistic way. Therefore, synergy is indexed by deviations from additivity.

#### Visualizing interactive effects

We developed an innovative way to unravel any existing interactions, borrowed from data science. The Individual Conditional Expectation (ICE) of an observation was estimated using the above equation 10 (i.e. 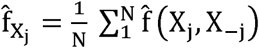), absent the averaging. We used plots of the ICE in order to disaggregate the estimated average marginal effect by displaying the estimated functional relationship for each observation. We therefore plot N estimated conditional expectation curves, each reflecting the individual predicted response as a function of the exposure X_j_, conditional on the observed X_−j_ (29). Unlike the marginal effects, this approach allowed us to observe and identify existing interactions.

We used non-parametric bootstrap (500 samples) for statistical inference for the estimated average causal effects (ACEs), interactive effects, and exposure–response relationships. In the absence of a theoretical formulae for the asymptotic distribution of these parameters within the SuperLearner framework, the bootstrap allows to approximate the 95% confidence intervals (CIs) (30, 31).

## RESULTS

### Predictive performance of the SuperLearner

The four simulated exposure-response relationships had different optimal *R*^2^ of 0.88, 0.67, 0.93, and 0.97, respectively for exposure-responses 1, 2, 3, and 4. The optimal value gives an upper bound on the possible *R*^2^ for each algorithm. Figure 2 shows the 10-fold cross validated MSE, the *R*^2^, as long as the *rel*MSE for each included algorithm for the four scenarios.

**Figure 2:**
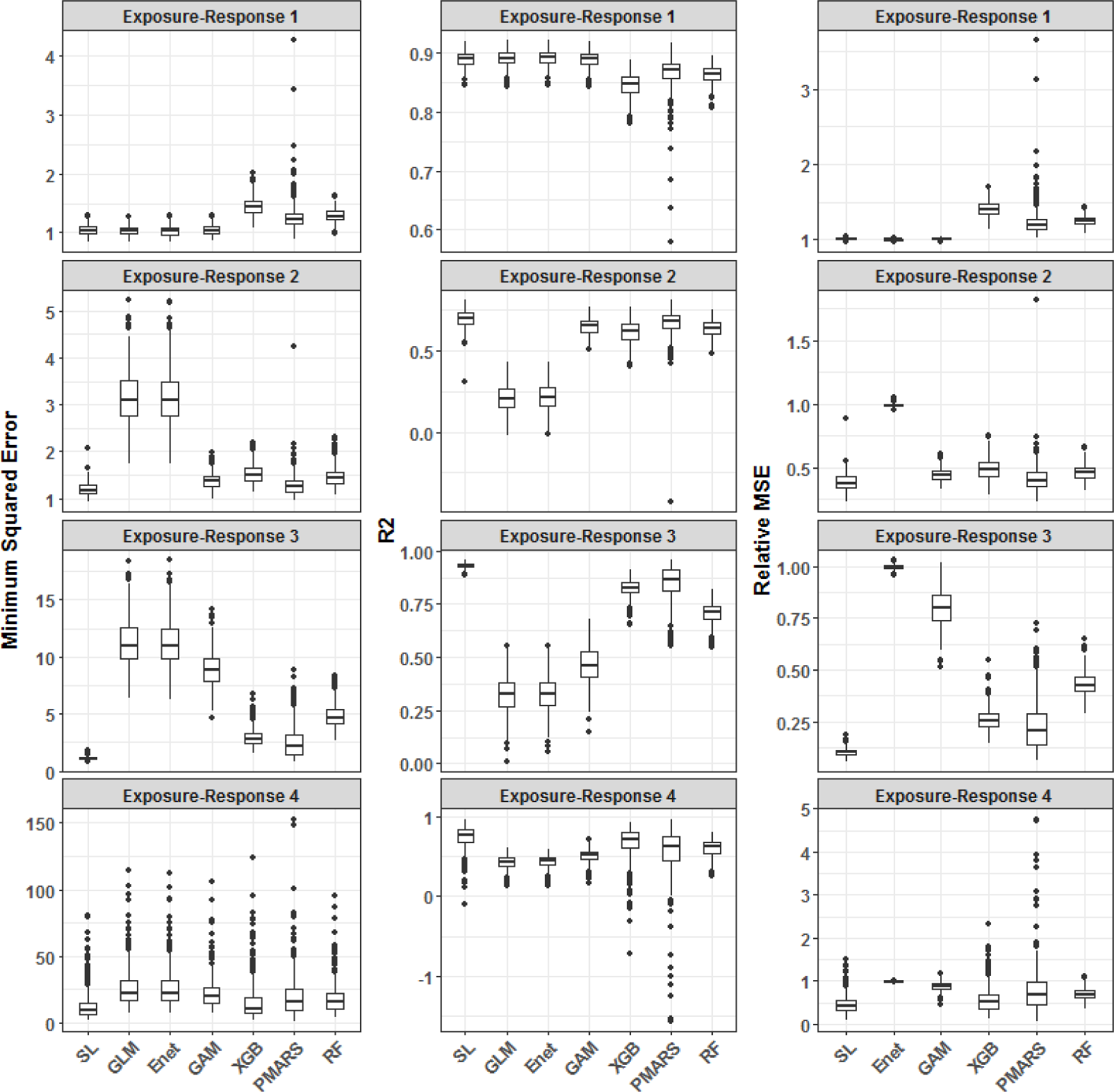
Distributions of the Minimum Squared error, R^2^, and *rel*MSE for the included algorithms, across the 4 scenarios. Regarding ACEs, Figure 3 shows the estimated ACEs for the 5 exposures for each algorithm and the 4 scenarios. Overall, the SuperLearner algorithm combined with G-computation was able to pick the true predictor in all the scenarios and did not pick any false positive reaching 100% sensitivity and specificity. The extreme gradient boosting algorithm (XGB) also reached the same level of sensitivity and specificity. None of the other algorithms performed as well as the SuperLearner and XGB algorithms. Regarding bias estimation, the SuperLearner with G-computation had the lowest average absolute bias, whereas the GLM had the highest average absolute bias (Results not shown).

**Figure 3:**
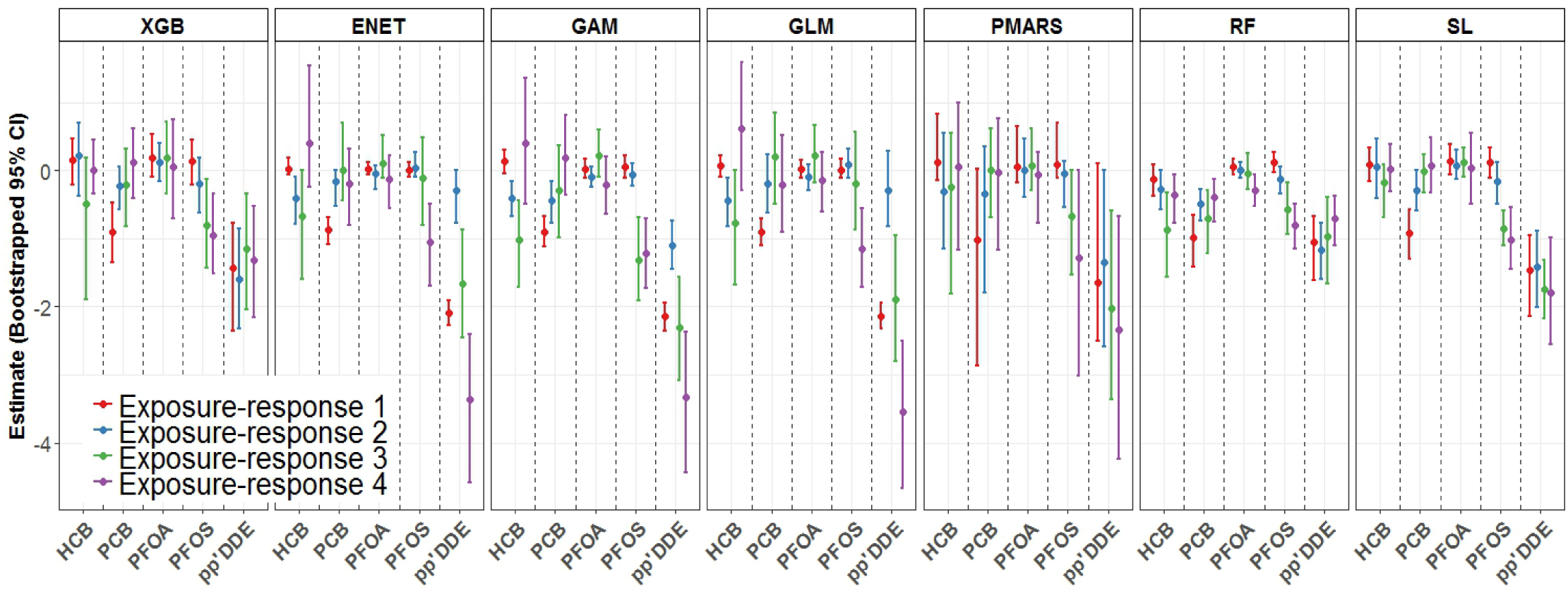
Average causal effects of the 5 exposures for each scenario, and for each included algorithm. *Reconstructing the dose response relationship:* We reconstructed the exposure-response relationship for each exposure and scenario (Figure 4). To accomplish this, we used the SuperLearner to predict the response at specific percentiles of the exposure of interest, and we reconstructed the dose-response by plotting these predictions along their bootstrapped 95% CI. Figure 4 shows that, using SuperLearner predictions, we were able to correctly reconstruct the exposure-response relationships for the four scenarios.

**Figure 4:**
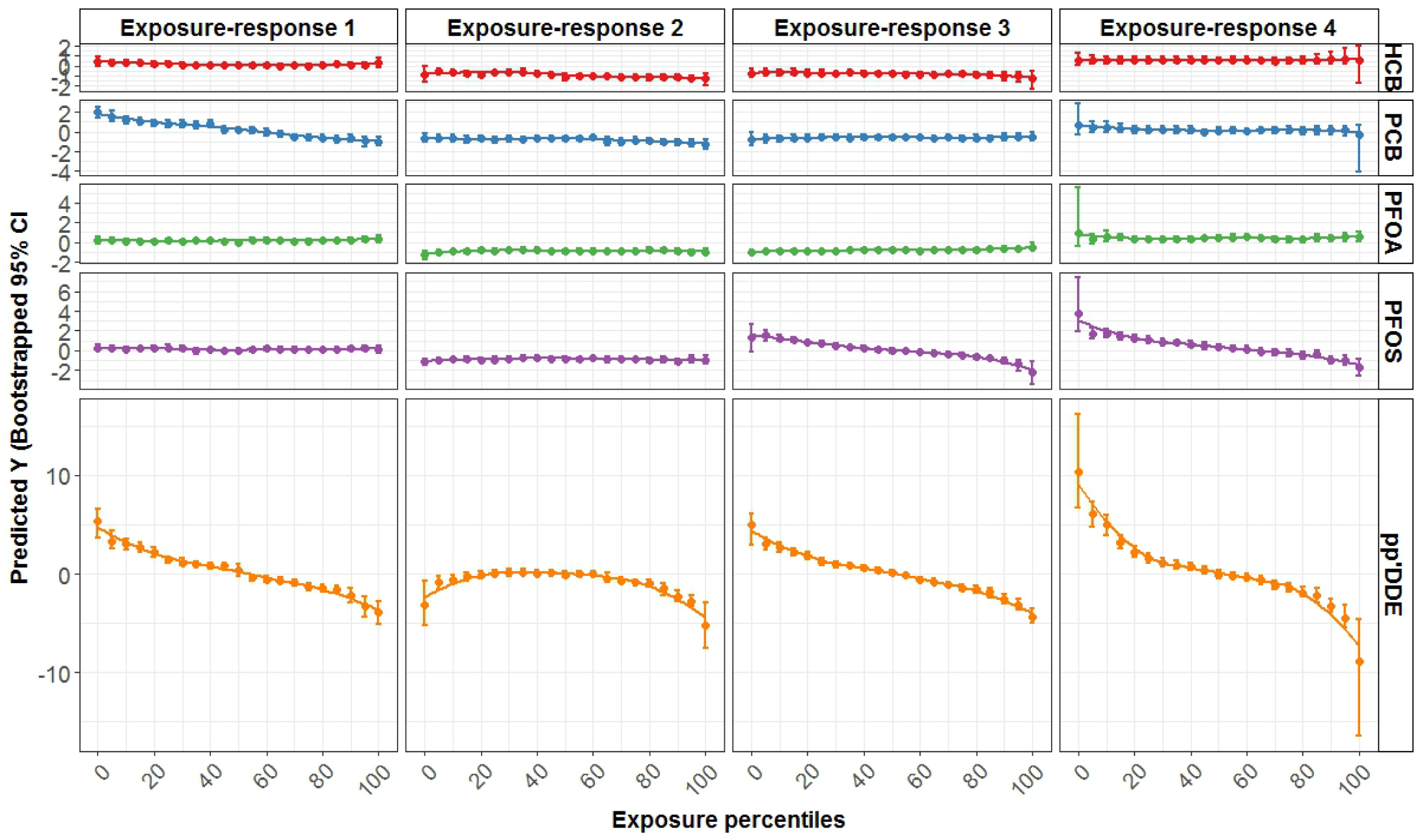
Exposure response relationships between the 5 exposures and the 4 generated outcomes. *Detection of interactions:* We calculated all potential two-way interactions using the predictions form the SuperLearner after plugging desired values of the two specific exposures of interest. The only interactive effects that are truly differed from 0 are the interactions between PFOS and pp-DDE for the simulated exposure-response 3, as well as the interaction between pp-DDE and sex for the simulated exposure-response 4. These were the only included interactions in the simulated data. Figure 5 shows interactive effect estimates for each pair of potentially interacting exposures, as well as sex. Effect estimates were derived from Equation 11 and 95% CI were constructed using bootstrap. Again, the SuperLearner was able to pick the two true interactions in Exposure-response 3 and 4. GLM, ENET, and GAM models were not able to pick interactions since these models do not handle interactions unless they are formally included in the model’s specification. XGB and PMARS algorithms performed very well although the former picked a false positive interaction in Exposure-response 1 and the later tended to be bounded by 0 when it picked interactions. RF algorithm tended to pick a high proportion of interactions, and was therefore keen to select false positive interactions. In regard to effect estimates, the SL algorithm was again the less biased one (Figure 5).

**Figure 5:**
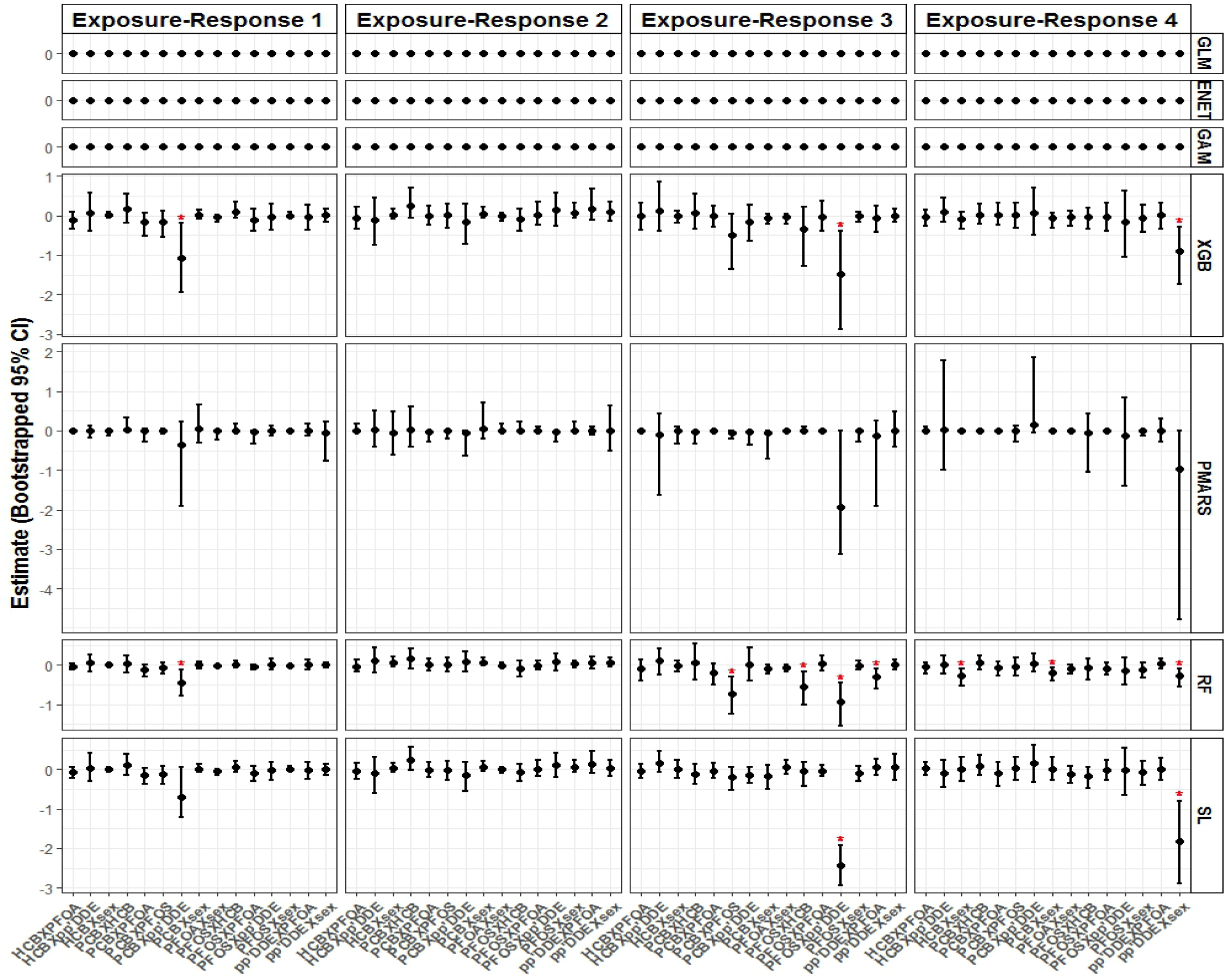
Effect estimates of the potential two-way interactions for all included algorithms and the 4 scenarios. Red asterisks indicate significant interaction terms. *Visualizing interactions:* In addition to this well-known approach embedded within the potential outcome framework, we present an innovative way to unravel existing interactions. We estimated the individual conditional expectations for each observation for specific levels of the exposure of interest (i.e. percentiles). Unlike Figure 4, where effects are averaged for all observations, plotting ICE allows observing specific patterns by groups of individuals (Figure 6).

**Figure 6:**
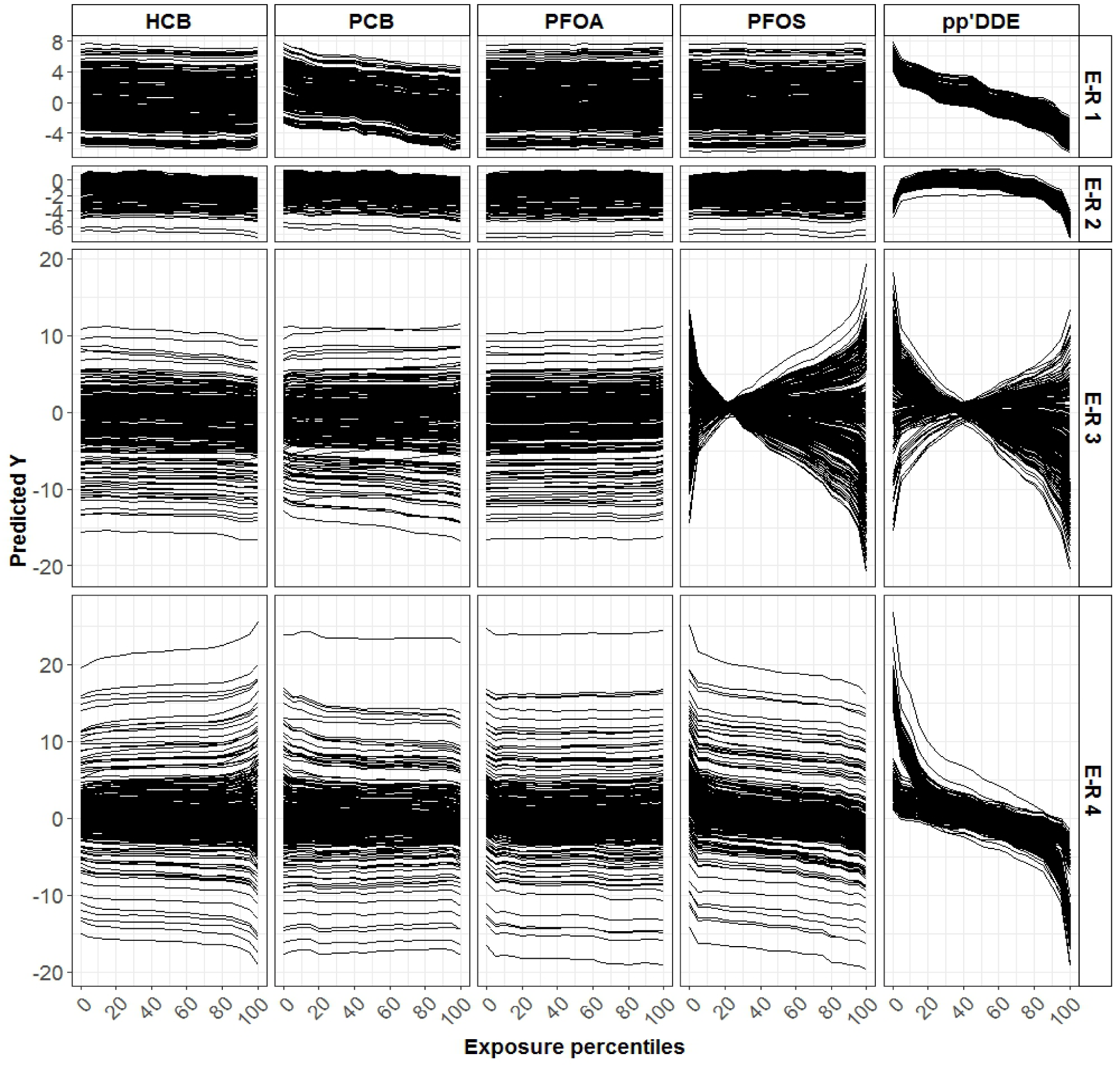
Individual conditional expectations for each exposure and for each generated exposure-response relationship. E-R: Exposure-Response. In this Figure 6, for most Exposure-Response relationships, ICE present parallel patterns for all the individuals, except when there is an interaction with another variable as demonstrated for *pp*’DDE in Exposure-Response 3 and 4, and for PFOS in Exposure-Response 3. Two examples are shown in Figures 7 and 8. Figure 7 describes the ICE for specific percentiles of *pp*’DDE exposure according to PFOS levels. Clearly, we can identify the interactive effect between *pp*’DDE and PFOS for Exposure-response 3 in Figure 7 since the dose-response relationship between *pp*’DDE and Y3 depends on the levels of PFOS.

**Figure 7:**
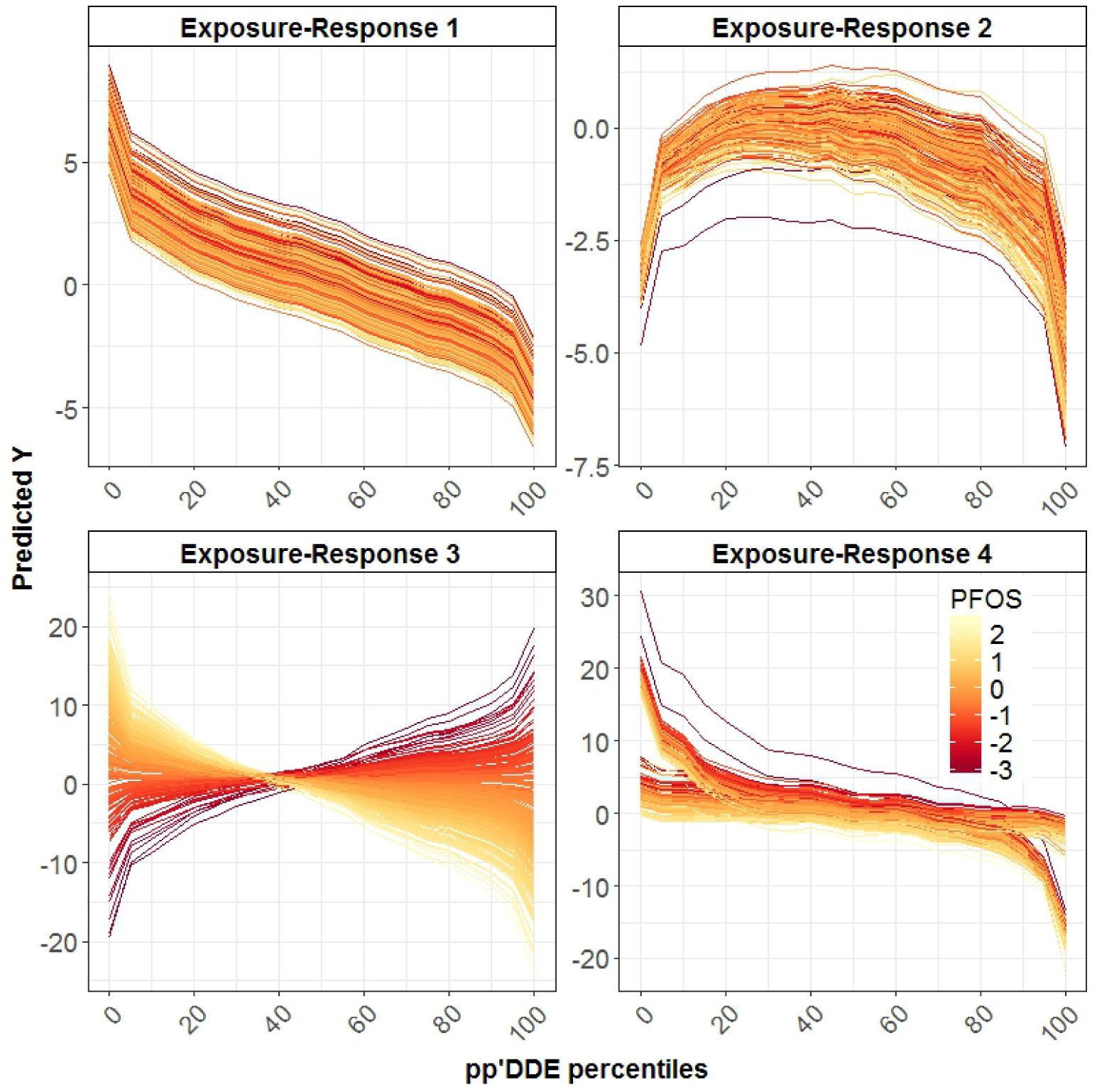
ICE of the relationship between *pp*’DDE and generated outcomes according to PFOS levels. Figure 8 describes ICE for specific percentiles of *pp*’DDE exposure according to sex. We can therefore visualize the effect modification by sex of the relationship between *pp*’DDE and Y4.

**Figure 8:**
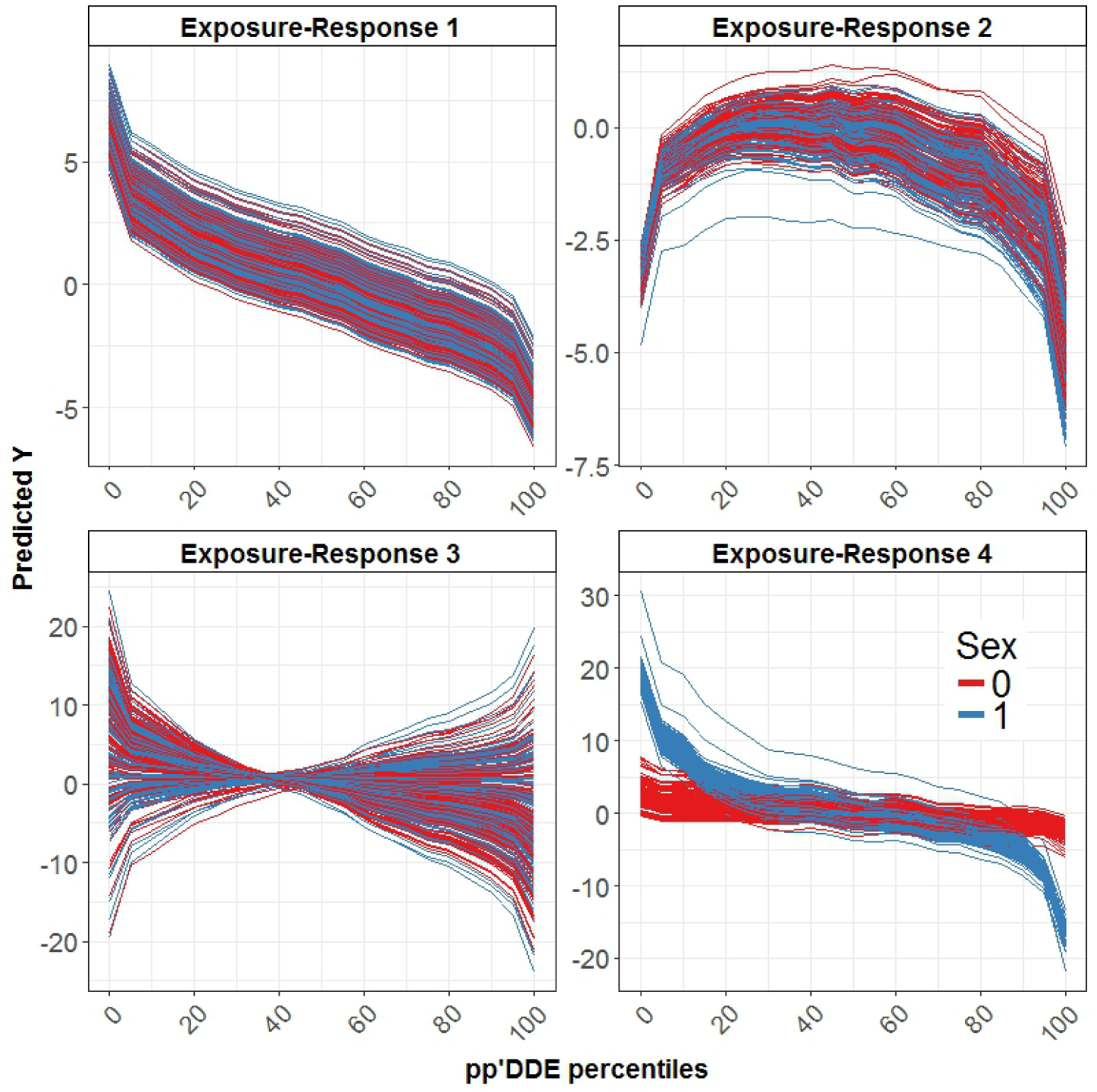
ICE of the relationship between *pp*’DDE and generated outcomes according to sex.

For the simulated exposure-response 1, most of the included algorithms performed very well given the simple linear relationship (Figure 2). Linear regression based methods perform best. Elastic net, GLM and the SuperLearner had the largest *R*^2^ (0.89). For exposure-responses 2, 3, and 4, the SuperLearner had the largest average *R*^2^ each time with *R s*^2^ of 0.69, 0.93, and 0.74. Results for the MSE exhibited the same pattern (Figure 2). Overall, the SuperLearner yielded the best average *R*^2^ (0.81) and the best average gain (lowest relMSE 0.48 in comparison to the widely used linear model across all the scenarios (Figure 2). Importantly, the SuperLearner was able to adapt to the true underlying structure of the data for each of the scenarios, and performed best or close to the best, demonstrating its ability to yield robust predictions across different scenarios.

### DISCUSSION

In the present paper, we have shown that ensemble learning methods to model the conditional mean of the outcome given known confounders can be advantageous, and have the potential to provide a solution to the long-lasting issue of chemical mixtures, especially in regard to model misspecification and multicollinearity issues. While this application represents a first proof of concept for application of these computational techniques to multiple correlated exposures in environmental epidemiology studies, our findings suggest that the proposed approach has an excellent predictive performance in a realistic environmental health scenario, in addition to a very good ability to correctly reconstruct dose-response relationships and to detect interactions. We were also able to derive estimates of marginal effects, using G-computation, a maximum likelihood substitution estimator. Under assumptions of conditional exchangeability, consistency, and positivity, these estimates may be interpreted causally.

We considered 4 simulations characterized by varying degrees of complexity to describe the Exposure-Response relationships. Each time, the SuperLearner combined with G-computation performed best or close to the best algorithm. This is an important and primordial result since the true parametric specification of the model is almost never known a priori. Therefore, the use of the SuperLearner allows relaxing the strong a priori assumptions in regard to the Dose-Response relationship.

The literature on the use of ensemble learning methods for estimating causal effects is limited (32), especially in the field of environmental epidemiology. To our knowledge, this approach is completely new and has not been explored in environmental epidemiological settings involving multi-pollutant exposures. Although ensemble learning techniques have gained increasing recognition, especially for climate change predictions (33-35), their use in environmental health studies is rare. Existing comparable attempts in this type are those of Diaz et al. (36), Chambaz et al. (37) and Kreif et al. (38), all of which are developed in contexts (e.g. genetics, health policy) very different from environmental mixtures. But these investigations did not extend the use of the SuperLearner to detect interactions and/or to construct dose-response relationships. Given that the marginal effects of an exposure in a nonlinear model are not constant over its entire range, even in the absence of interaction terms, it is important to reconstruct the dose response relationship (39).

Several methods have been proposed to estimate the joint effects of environmental mixtures, and individual effects within a mixture, often with an emphasis on variable selection. The most widely used methods are the LASSO (40), EWAS (5, 41), weighted quantile sum regression (42, 43), and Elastic Net (13, 44). A major disadvantage of such approaches is that they typically assume specific and often restrictive parametric functional forms for the exposure-response relationship, often resulting in a model that does not accurately capture the complexity of the relationships among high dimensional covariates and health outcomes. This misspecification can lead to biased estimators and overly liberal (too optimistic) assessment of the uncertainty associated with estimation. We typically observed this trend in our results when such methods were applied to complex Exposure-responses with interactions and non-linearities.

The present work has several limitations. First, we only considered environmental exposures and one potential effect modifier, eg. sex. Future studies should examine the performance of this approach while taking into account the data generating process of such exposures allowing for multiple potential confounders. The inclusion of potential confounders when these are measured, should be based on a priori knowledge and causal approaches based on the interdependencies between variables, e.g. Non parametric structural equation models (45, 46), and not only based on the predictive performance. Second, the ability of the SuperLearner approach depends on the choice of candidate learners that should be guided by theoretical and practical considerations based on expert knowledge. We did not include some powerful algorithms such as the Bayesian Additive Regression Tress (BART), nor did we specify tuning parameters for the included algorithms to minimize computational time. But these are possibilities that should be investigated in future work. Third, we used the bootstrap to estimate valid confidence intervals in the absence of a theoretical formula for the asymptotic distribution of the parameters of interest within the SuperLearner framework. This gave rise to heavy computational burden, since the described analyses are quite time consuming. Running these analyses in multicore parallel computing will substantially reduce this time, but just how much time can be saved depends on the availability of computing cores with sufficient memory, and is therefore installation-dependent (47). Finally, the generated dose-response relationships in this investigation were based on associations with relatively strong effect sizes, and further studies will consider the performance of such approach when effect sizes are moderate or low.

Despite the abovementioned limitations, these analyses provide a significant contribution to the field of environmental health. This approach leverages the high predictive ability of ensemble learning techniques, while opening the blackbox of these methods to allow for the estimation of individual associations, interactive effects, and reconstruction of dose-response relationships. The overall idea of this paper is to propose a general approach that is flexible. Such method can lay the ground for additional methodological extensions allowing for future developments encompassing high dimensional data from omics technologies such as microbiomics, epigenetics, and metabolomics, as well as a flexible way to assess mediation and moderation, as well as multivariate analyses. This approach can also be easily adapted to estimate generalized propensity scores for doubly robust estimations. Therefore, additional developments and progresses starting from this work may provide substantial improvements.

